# Learning Distance-Dependent Motif Interactions: An Explicitly Interpretable Neural Model of Genomic Events

**DOI:** 10.1101/2020.08.27.270967

**Authors:** Thomas P. Quinn, Dang Nguyen, Phuoc Nguyen, Sunil Gupta, Svetha Venkatesh

**Affiliations:** Applied Artificial Intelligence Institute (A2I2), Deakin University, Geelong, Australia

## Abstract

In many biological studies, prediction is used primarily to validate the model; the real quest is to understand the underlying phenomenon. Therefore, interpretable deep models for biological studies are required. Here, we propose the **Hyper**-parameter e**X**plainable Motif **Pair** framework (**HyperXPair**) to model biological motifs and their distance-dependent context through explicitly interpretable parameters. This makes **HyperXPair** more than a decision-support tool; it is also a hypothesis-generating tool designed to advance knowledge in the field. We demonstrate the utility of our model by learning distance-dependent motif interactions for two biological problems: transcription initiation and RNA splicing.

## 1 Introduction

Living cells store information in large repeating chains of molecules called polymers. Example polymers include DNA, RNA, and proteins, each of which can be described as a sequence composed from a finite set of bases. The function of any molecule ultimately depends on the chemical properties of its atomic constituents. Yet, it is possible to make accurate predictions directly from 1D sequence representations through biologically meaningful sub-sequences called *motifs*. Identifying motifs is critical to understanding the Protein-DNA and Protein-RNA interactions involved in cancer and its treatment Bonnal et al. [2020].

Interest has gathered recently around how machine learning, especially deep convolutional neural networks (CNNs), can predict genomic events like RNA splicing [Ching et al., 2018, Jaganathan et al., 2019, Wang et al., 2019, Albaradei et al., 2020]. Although deep CNNs achieve good performance, they do not learn an explicitly interpretable model. *For many biological studies, prediction itself is used primarily to validate the model; the real quest is to understand the underlying phenomenon. Therefore, interpretable deep models for biological studies are required*.

Learning an interpretable model of genomic events requires solving what we call the *distance-dependent motif interaction problem* (presented in Figure 1). This problem poses 2 key challenges:

- **Motif degeneration:** Biological motifs are ill-defined constructs, and usually have fuzzy definitions. For example, GGAG and AGGA are both, among others, valid Shine-Dalgarno sequences [Ma et al., 2002].
- **Distance-dependence:** A motif may be necessary, but not sufficient. Whether a motif is sufficient depends on its context, such as its distance from another motif. In other words, motif pairs act synergistically, not additively [Ke and Chasin, 2010], as a function of their proximity. In the case of transcription initiation, the observed frequencies of inter-motif distances follow a Gaussian distribution [Ma et al., 2002].

**Figure 1:**
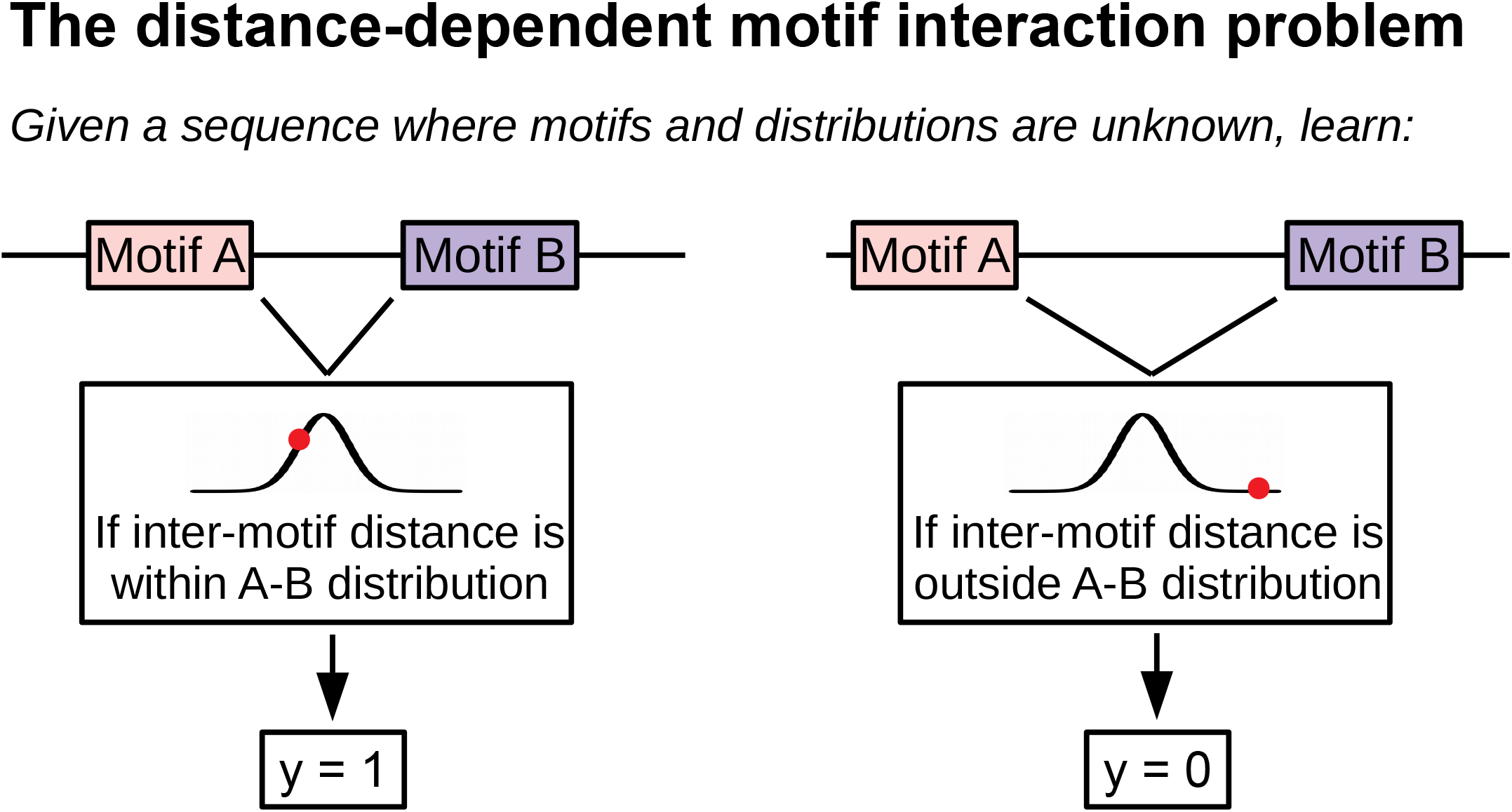
When Motifs A and B are both present *and* their distance falls within the inter-motif distance distribution, then a genomic event (represented by *y* ∈{0, 1}) will occur. Our model learns the motif identities and inter-motif distance distributions explicitly.

We propose the **Hyper**-parameter e**X**plainable Motif **Pair** framework (**HyperXPair**) as an interpretable “clear box” solution to the distance-dependent motif interaction problem. It has 2 modules whose parameters explicitly represent the aspects we want to interpret:

- **Motif identity module:** This is simply a 1-layer CNN where the kernel parameters represent the consensus motifs. This module solves the challenge of motif degeneration because the CNN kernel can capture several overlapping motifs. In the case that motif degeneration is too extreme, separate filters could capture alternate versions.
- **Motif distance module:** This is also a 1-layer CNN, but with a custom kernel. The kernel parameters depend on *learnable hyper-parameters that represent inter-motif distances as a distribution*, which we can efficiently search for using Bayesian optimization [Shahriari et al., 2016]. This module solves the challenge of distance-dependence because the CNN kernel can weigh motif interactions as a function of the distance between them.

Unlike other models, **HyperXPair** learns to relate biological sequences to genomic events through parameters that are immediately understood by a biologist. This makes **HyperXPair** more than a decision-support tool; it is also a hypothesis-generating tool designed to advance knowledge in the field. We demonstrate our model by learning distance-dependent motif interactions for two biological problems: transcription initiation and RNA splicing. We show how an interpretation of the model’s parameters can be combined with literature to extend domain knowledge with new hypotheses about genome biology.

## 2 Related work

### Distance-dependent motif interactions

We hypothesize that two key factors determine whether a motif will execute a genomic event. The first is *motif identity*: certain sub-sequences within the DNA are required to execute an event. They are short (1s-10s of bases long) and fuzzy. The second is *motif context*: motifs are necessary, but not always sufficient. Events may require multiple motifs separated by a specific number of positions. We base this hypothesis on the genome biology literature. For example, consider transcription initiation. The Shine-Dalgarno sequence needs to be ∼5-13 bases upstream from the Kozak consensus sequence to initiate transcription [Ma et al., 2002]. Another example is RNA splicing, where RNA has sub-sequences removed and concatenated [Ule and Blencowe, 2019] (e.g., to convert “machine” to “m…ine” to “mine”). Here, fuzzy motifs cooperate in a distance-dependent fashion, potentially involving long-range interactions [Wong et al., 2016].

### Explainable AI for biological sequences

There are two general approaches to make “black-box” neural networks more comprehensible: (1) to train an arbitrary neural network and infer predictions secondarily via *post-hoc analysis*, or (2) to train a neural network whose parameters can be interpreted directly. Among post-hoc methods, Zhou and Troyanskaya [2015] have used a perturbation method called *in silico mutagenesis* to identify motifs by “mutating” DNA bases and measuring the change in prediction. Intuitively, important bases will cause bigger prediction changes when perturbed. Koo et al. [2018] used a similar procedure to understand how motif number and spacing impacts prediction. Meanwhile, a back-propagation method called DeepLIFT [Shrikumar et al., 2019] identifies motifs by comparing the gradient for a sample-of-interest against a reference [Wang et al., 2019]. In contrast, our model takes inspiration from *self-explanation*, in which interpretability is built-in architecturally [Alvarez-Melis and Jaakkola, 2018]. In **HyperX-Pair**, the parameters explicitly represent the aspects we want to interpret.

### RNNs for biological sequences

The 1-dimensional spatial arrangement of a biological sequence lends itself to analysis by recurrent neural networks (RNNs), which can detect sequential patterns with some tolerance to changes in the position and identity of the motifs [Hawkins and Bodén, 2005]. Long short-term memory (LSTM) is most popular today, owing to its ability to learn patterns with long time lags [Hochreiter and Schmidhuber, 1997]. LSTM has been applied to biological sequences, often in combination with CNNs. For example, Hassanzadeh and Wang [2016] and Pan et al. [2018] both propose a two-tier model where an initial CNN layer learns to represent DNA motifs, while a second LSTM layer learns long-range interactions between motifs. We include LSTMs in our baselines.

### CNNs for biological sequences

Encoding the DNA as a matrix makes it image-like. If we conceptualize a motif as a visual stimulus, then we can understand why CNNs are a natural choice for learning translation-invariant motifs [Fukushima, 1980, Waibel et al., 1989]. Convolutional filters from the lowest layers tend to learn motif fragments, which get assembled in deeper layers to capture higher-order interactions and distance-dependence. Deep CNNs have enabled many sequence prediction tasks, from splice prediction [Jaganathan et al., 2019] to drug-target affinity prediction [Nguyen et al., 2020a]. One potential advantage of deep CNNs is that they do not require prior knowledge about how an event happens [Bao et al., 2019]. We include CNNs in our baselines.

### Interpretable CNNs

Biological sequences are different than images. For images, lower-layer features like edges may not mean much to the investigator; for biological sequences, lower-layer features are the motifs themselves, and thus should be preserved accurately [Koo and Eddy, 2019]. Unfortunately, deep CNNs can cause the lower layers to fragment into partial motifs that misrepresent the underlying biological reality [Koo and Eddy, 2019]. Several studies have explored how architectural variants could encourage lower layers to learn coherent motifs, such as exponential activation [Koo and Ploenzke, 2019], Gaussian noise injection [Koo et al., 2019], or kernel regularization and constraints [Ploenzke and Irizarry, 2018]. However, even if a network could learn complete motifs, the interactions may get obscured in the depths of the CNN [Koo and Eddy, 2019]. Our architecture learns interactions explicitly.

## 3 Model Overview

Figure 2 and **Supplemental Figure 1** illustrate the 3 stages of the **HyperXPair** architecture. First, a CNN learns motif identities from a 1-hot embedding of DNA, producing a **motif activity map** that describes the presence of each motif along the sequence. Second, another CNN uses a **motif distance kernel** to compute interaction scores for each motif pair along the sequence. Third, a global max pool layer selects the largest interaction score for each pair, and uses them to predict the outcome via a simple regression.

**Figure 2:**
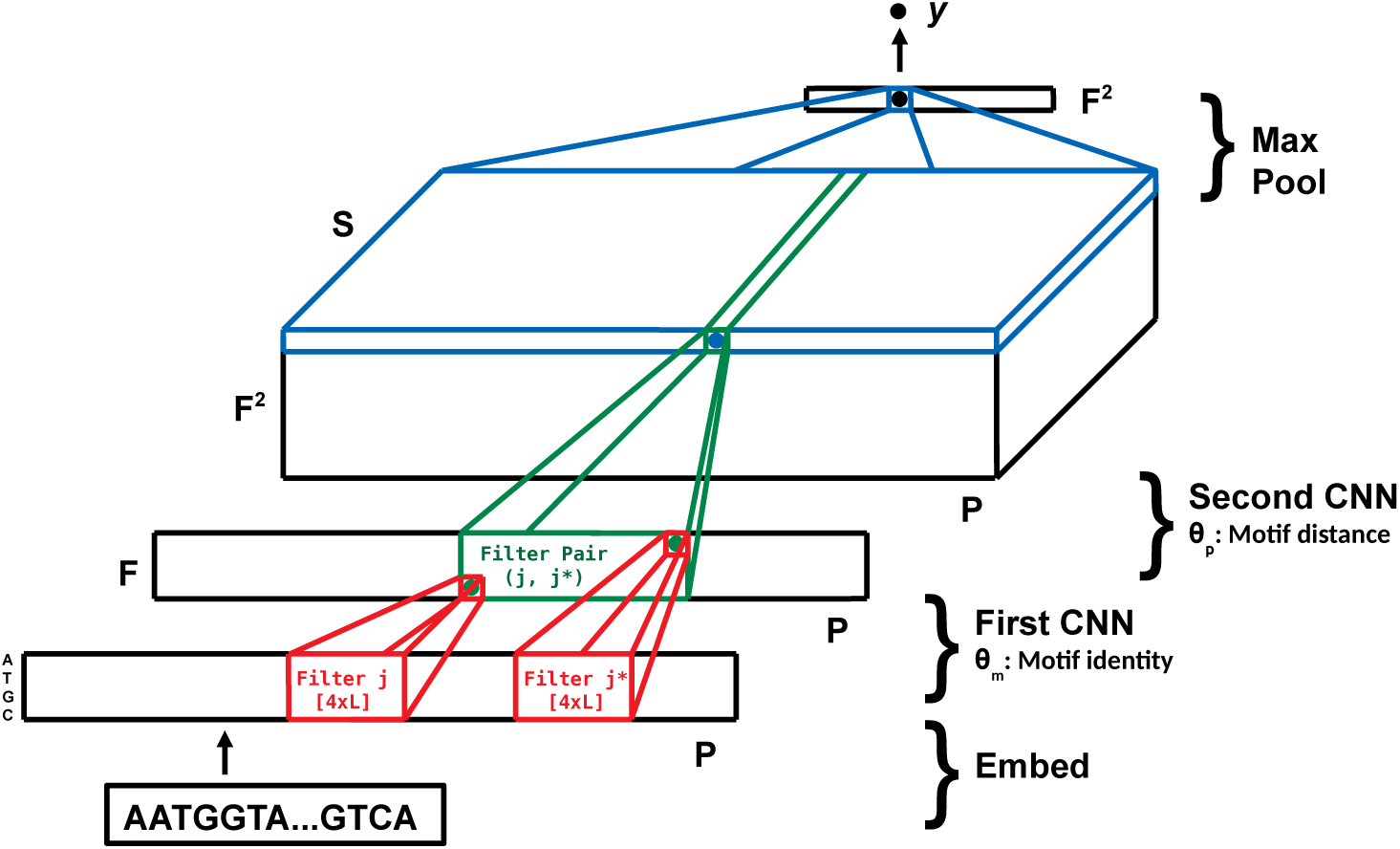
**HyperXPair** works in 3 stages: (1) a first CNN learns motifs from an embedded sequence, (2) a second CNN computes distance-dependent interaction scores, and (3) a global max pool layer predicts the output.

### 3.1 Stage 1: First CNN Discovers Consensus Motifs

Before we can make predictions based on the distances between motifs, we need to learn the motifs themselves. The first function converts the input **x** into a **motif activity map** describing motif presence along the sequence:

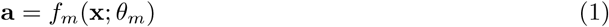

where *f*_*m*_ is a CNN and *θ*_*m*_ has *F* filters sized *L*-by-4, where *L* is the maximum motif length. **Interpretation of module:** *Each filter represents a motif that can be normalized by a weighted softmax transform and visualized directly by a sequence logo plot* Schneider and Stephens [1990].

Note the input **x** contains *N* genomic sequences, having up to *P* bases with 4 possible states. We 1-hot encode the 4 states–“A”, “T”, “G”, or “C”–then add *P* bases of “padding” to both sides of the sequence. This results in a tensor of size [*N*× 3*P*× 4]. The padding allows the model to capture interactions at the ends of the sequences.

### 3.2 Stage 2: Second CNN Considers Distances

Now that we have a **motif activity map**, we can score motif interaction based on the distance between pairs. For *F* motifs, there are *F* ^2^ motif pairs. Our goal is to compute a distance-dependent interaction score for each pair *j* and *j*^*^. The score should be large when the motifs are present *and* their distance falls within an assumed inter-motif distance distribution. The score should be small when a motif is absent *or* their distance falls outside the distribution.

We describe below how we efficiently search for the parameters of the distance distribution using Bayesian optimization. For now, let us assume we already know the mean *M* ^(*j,j*\*)^ and standard deviation Σ^(*j,j*\*)^ that define a Gaussian distance distribution, forming the basis of a **receptive field** that assigns weight to motif *j*^*^ as a function of its distance from motif *j*. The receptive field scores each motif gap {0…*S*−1} where *S*−1 is the maximum motif gap.

Figure 3 shows how we can encode the **receptive field** as a custom CNN kernel that scores interaction as a function of distance. The **motif distance kernel** will have the size [*F* ×*S*×*S*], where the first slice of the kernel captures a motif gap of 0 (i.e., when motif *j* and *j*^*^ overlap), the second slice of the kernel captures a motif gap of 1 (i.e., when motif *j* and *j*^*^ are separated by one position), and so on. The kernel 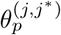 thus follows deterministically from the receptive field:

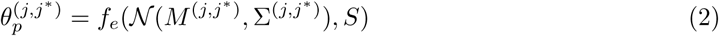

where *f*_*e*_ is the encoding rule. We do not apply the receptive field directly because it would weigh multiples of one filter the same as a pair. Empirically, this does not work well.

**Figure 3:**
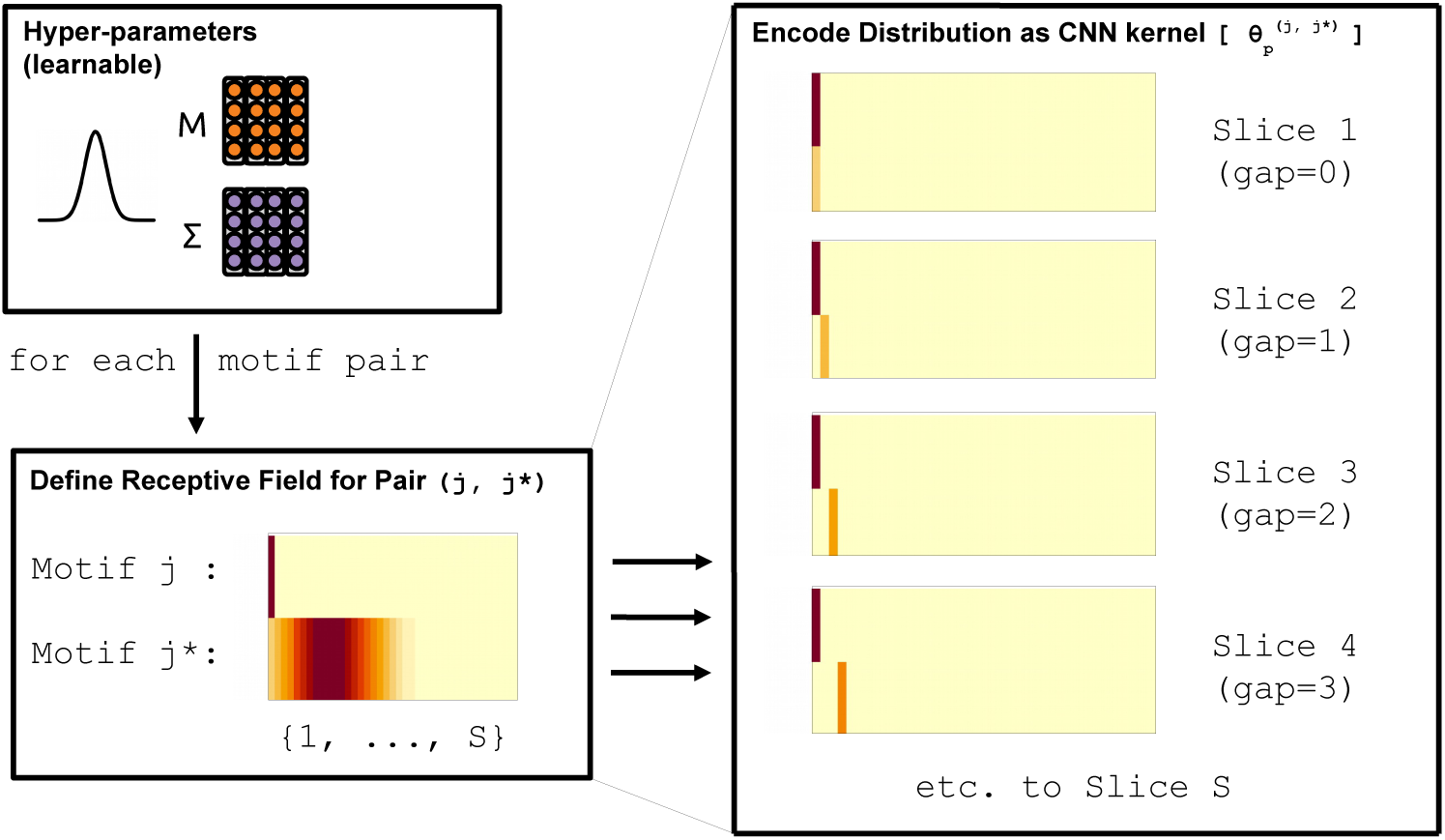
The hyper-parameters *M* ^(*j,j*\*)^ and Σ^(*j,j*\*)^ define the **inter-motif distance distribution** as a receptive field that assigns weight to motif *j*^*^ as a function of its distance from motif *j*. When a motif pair falls within the red bands, it is within the distance distribution and will receive a high score. When a motif pair falls within the yellow-orange bands, it is outside the distance distribution and will receive a lower score (with the score approaching zero for the brightest hues of yellow).

Note that *S* gives *M* ^(*j,j*\*)^ contextual meaning. When *M* ^(*j,j*\*)^ = 0, a motif gap of *S/*2 receives the highest weight. When *M* ^(*j,j*\*)^≈−3, a gap of 0 receives the highest weight. When *M* ^(*j,j*\*)^≈ 3, a gap of *S* receives the highest weight. The Σ^(*j,j*\*)^ determines how the weights decrease as the observed gap gets further from the optimal gap. **Interpretation of module:** *These hyperparameters define the mean and standard deviation of a receptive field that can be visualized directly as the inter-motif distance distribution*.

We apply the kernel directly:

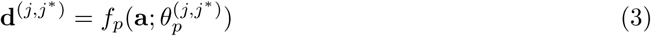

where *f*_*p*_ is a CNN and **a** is the motif activity map from above. The resultant tensor **d**^(*j,j*\*)^ represents the distance-dependent interaction score for each motif gap {0…*S*−1} at each DNA position {1…*P*}.

A high interaction score implies that both motifs are present *and* their distance falls within the distribution, thus solving the distance-dependent motif interaction problem.

### 3.3 Stage 3: Interaction Scores Regressed to Output

We calculate the max interaction score for each motif pair as the global maximum of **d**(*j,j*\*):

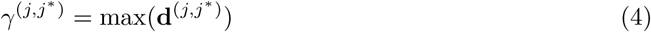

This results in a single number for each motif pair. When repeated for all *F* ^2^ motif pairs, we get a tensor of size [*N* × *F* × *F*], or, equivalently, [*N* × *F* ^2^]. Thus,

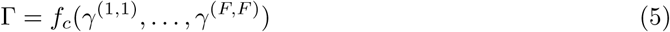

where *f*_*c*_ is a concatenation. We can predict the outcome:

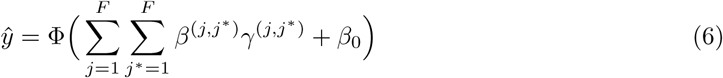

where *β*^(*j,j*\*)^ are the linear model coefficients for a motif pair *j* and *j*^*^, and Φ is the generalizing link function. Here, our outcome is binary, so Φ is a sigmoid transform.

Although Eq. (6) represents a direct connection to the output, nothing stops us from adding more layers if it were appropriate for the dataset size and/or problem complexity. Our novel *motif distance module* could occur before a deep multilayer perceptron (and/or after a deep CNN), allowing the network to be very deep. We restrict the total number of layers for two reasons: (1) to bring more interpretability to the motif identity module, and (2) to not distract from our key contribution: the motif distance module.

## 4 Model Learning

Learning occurs in two loops. In the inner-loop, we use stochastic gradient descent to search for the motifs whose distance-dependent interactions minimize training loss. In the outer-loop, we use Bayesian optimization (**BO**) to search for the hyper-parameters {*M*, Σ}that give the best neural network. Algorithm 1 describes our implementation.

**Algorithm 1:**
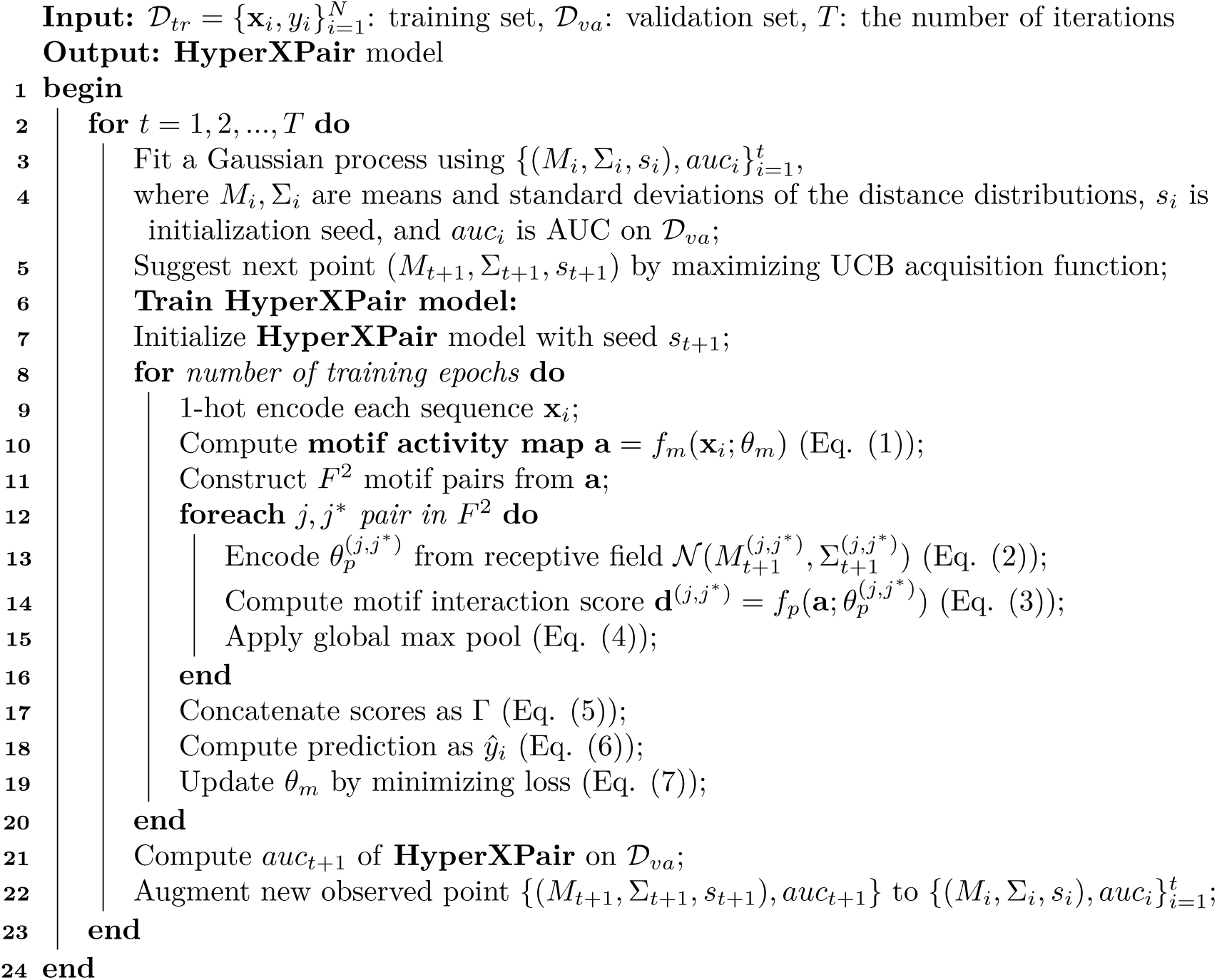
The proposed **HyperXPair** algorithm.

### 4.1 Inner-loop: motif discovery

For a given set of hyper-parameters, we train our neural network end-to-end to minimize the loss

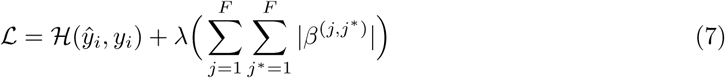

where *ℋ* is binary cross-entropy, and *λ* penalizes the magnitude of the weights in the final layer. This regularization allows us to over-specify the number of motifs needed for a model, which may be unknown.

During early experiments, we noticed two drawbacks with the vanilla architecture. First, it quickly overfitted to the training data by learning elaborate motifs not found in the validation set. We address this by adding an aggressive dropout layer of 30% to the embedded sequence input. Second, it would sometimes fail to converge. We address this by initializing the model across several random seeds, and choosing the best performer *based on the validation set*.

### 4.2 Outer-loop: inter-motif distance discovery

We optimize 3 hyper-parameters: the means and standard deviations of the distance distributions, and the initialization. To choose them, we use BO [Snoek et al., 2012, Nguyen et al., 2020b], implemented in rBayesianOptimization. Here, we treat the classification process of our model as a black-box function, where the inputs are the means, standard deviations, and initialization seed, and the output is validation set AUC. We then use a Gaussian process (GP) [Rasmussen, 2003] with an Upper Bound Confidence (UCB) acquisition function [Srinivas et al., 2012, Snoek et al., 2012].

To speed up optimization, we only consider a half-matrix of *M* ^(*j,j*\*)^ where *j* < *j*^*^, and further assume that all Σ^(*j,j*\*)^ have the same value. This greatly reduces the number of variables we need to learn. The decision to treat initialization as another variable is pragmatic, based on early experiments where some initializations more readily produced relevant motifs. In our implementation, the validation set is re-sampled, and the network weights re-learned, at each BO iteration.

## 5 Experiments

### 5.1 Data and baselines

#### 5.1.1 Synthetic data

We simulated data based on the Stormo et al. [1982] case study. Using a custom script, we simulated DNA from 2 classes. In the first class, the DNA contained the GGAGG motif ahead of the ATG motif by ∼ 5-13 positions (with the actual distance sampled from a Gaussian distribution where *µ* = 8 and *Σ* = 2, as approximated from [Ma et al., 2002]). In the second class, the DNA either (a) contained the GGAGG motif *after* the ATG motif, or (b) did not contain the ATG motif.

We simulated *N* = [128, 256, 512, 1024, 2048, 4096] total samples so that we could test our model over a range of sample sizes. We repeated the simulation procedure 5 times, withholding 33% of samples as a test set (yielding 30 test sets overall). Note that because of nested validation, *N* = 1024 would imply that **HyperXPair** is trained on just 439 samples, with 109 for monitoring neural network validation loss, 137 for BO validation, and the rest for testing.

#### 5.1.2 Real data

Albaradei et al. [2020] benchmarked a deep CNN model for splice prediction, and made their data publicly available. We downloaded the *D. melanogaster* data^1^, chosen because fruit fly splicing is well-studied [Mount et al., 1992]. The data include two binary classification problems: (1) “donor” splice site present vs. absent, and (2) “acceptor” splice site present vs. absent. Each sequence contains 602 bases, centered on the splice site. We performed 5 separate random splits of the data, withholding 33% of the data each time (yielding 10 test sets overall).

#### 5.1.3 Baselines

We compare **HyperXPair** against several baselines:

- **Natural Language Processing (NLP) type methods**: By treating DNA as a document, and bases as words, we can represent DNA via the conventional NLP methods bag-of-words (**BOW**) and term frequency-inverse document frequency (**TF-IDF**) [Arora et al., 2018]. After constructing feature vectors, we train a linear SVM, tuning the hyper-parameter *C ∈*{0.001, 0.01, 0.1, 1, 10, 100, 1000} with a validation set. *Justification for baseline(s):* They perform well for standard NLP tasks [Arora et al., 2018].
- **Sequence embedding methods**: We also compare with **Sqn2Vec**, a neural embedding method for learning sequence representations [Nguyen et al., 2018]. This method learns a sequence embedding by predicting frequent sub-sequences (e.g., motifs) that belong to a sequence. Sqn2Vec has two models: Sqn2Vec-SEP and Sqn2Vec-SIM. *Justification for baseline(s):* They perform well for sequential data that have a small vocabulary, like DNA.
- **2-filter CNNs:** We implement a basic CNN with a 1D convolutional layer containing 2 filters of size 4×8, followed by a global max pooling layer connected to the output (**F2-CNN**). We also implement a variant with an intermediate 64-node dense hidden layer (**F2-CNN-Dense**). *Justification for baseline(s):* They use the same training specifications as **HyperXPair**, allowing us to isolate the contribution of the motif distance module from other factors.
- **Shallow CNNs:** We also implement a wider shallow CNN having a 1D convolutional layer with ReLu activation. As above, a global max pool layer that predicts the output. For these CNNs, *F* is set to{15, 20, 25, 30, 35, 40} for {128, 256, 512, 1024, 2048, ≥4096} samples. See “Deep CNNs” below for training specifications. *Justification for baseline(s):* They resemble the more interpretable models used for biology prediction tasks.
- **Deep CNNs:** The deep CNN begins the same as the shallow CNN, except that the global max pool layer is replaced by a [3×1] max pool layer. It then has two more sets of convolutional and max pool layers (using 2*F* and 4*F* filters, respectively), followed by a global max pool layer. It also has a 100-node dense hidden layer before the final output layer. Here, *F* is set to{5, 10, 15, 20, 25, 30} for {128, 256, 512, 1024, 2048, ≥4096} samples. All hidden layers have ReLu activations and the output layer has sigmoid activation. For training, we use the cross-entropy loss and ADAM optimizer with a learning rate of 0.001, batch size of 16, and 100 epochs. *Justification for baseline(s):* They resemble the high-performing models used for biology prediction tasks.
- **LSTMs:** For our LSTM, the dimension of symbol embedding is 128, the number of LSTM hidden units is 100, and the drop-out rate after each layer is 0.2. For training, we use the ADAM optimizer with a learning rate of 0.001, batch size of 64, and 50 epochs. *Justification for baseline(s):* They resemble the high-performing models used for biology prediction tasks.

#### 5.1.4 HyperXPair training

Unless otherwise noted, we used *F*≥ 2 total filters (chosen to simplify interpretation and improve run-time), *L* = 8 motif length (chosen because we are interested in short motifs), and *S* = 40 motif gap (chosen because we are interested in proximal motifs). If *F* > 2, *λ* = 0.01, else *λ* = 0.001. For training, the ADAM optimizer is used with a learning rate of 0.001 and a batch size of min(*N/*64, 320) (this lets all epochs have a similar number of batches). Training occurs for 50 epochs during hyper-parameter tuning, or 150 epochs otherwise (chosen based on early experiments with toy data). In all cases, early stopping occurs if validation loss does not improve for 15 epochs.

For BO, we run 80 iterations after a random initialization of 8 trials (based on a “rule-of-thumb” of 2 initializations and 20 iterations per variable being optimized for a single pair). In addition to tuning all *M* (bounded [−3, 3]) and one Σ (bounded [0.1, 3]), we tune the initialization seed over 1…10 (if *F* = 2) or 1…15 (if *F* > 2).

## 6 Results

### 6.1 All model parameters are explicitly interpretable

The model parameters describe: (a) the identity of the motifs, (b) the inter-motif distance distribution for each motif pair, and (c) the regression weights of each motif interaction as it relates to the final outcome.

- **Motif identities:** The filter weights from the first CNN layer define the motifs. They are interpretable because large positive weights imply that a DNA base is often present, while large negative weights imply that a DNA base is rarely present. A row-wise weighted softmax transform normalizes the filter weights into a position weight matrix (PWM) that can be visualized by a sequence logo plot Schneider and Stephens [1990]. (The softmax weight acts like a “gain knob” to magnify the signal-to-noise ratio of the associated information content, making visualization easier.)
- **Motif interactions:** The filter weights from the second CNN layer describe how the importance of an interaction relates to inter-motif distance. They are interpretable because they can be plotted to visualize the inter-motif distance distributions directly.
- **Regression weights:** The weights from the final regression layer are multiplied against the max interaction scores to predict the outcome. They are interpretable because the weights – arising from a linear model – describe the relative importance of the motif interaction inputs.

### 6.2 Study 1: Simulated transcription initiation data

Simulated data allow us to test whether our model can learn the correct motifs and inter-motif distance distributions. We consider two lines of evidence: (1) model performance, and (2) validity of learned motifs and distance distributions.

Figure 4 shows excellent performance for **HyperXPair** as compared with several baselines. Figure 5 shows how **HyperXPair** has correctly learned that having the motif GAGG∼5-13 bases ahead of ATG will predict transcription initiation. This figure explains how the model works, and is produced directly from the model’s parameters. **Supplemental Figures 2-6** show all filters for the 5 separate *N* = 4096 training sets, which agree with the one here.

**Figure 4:**
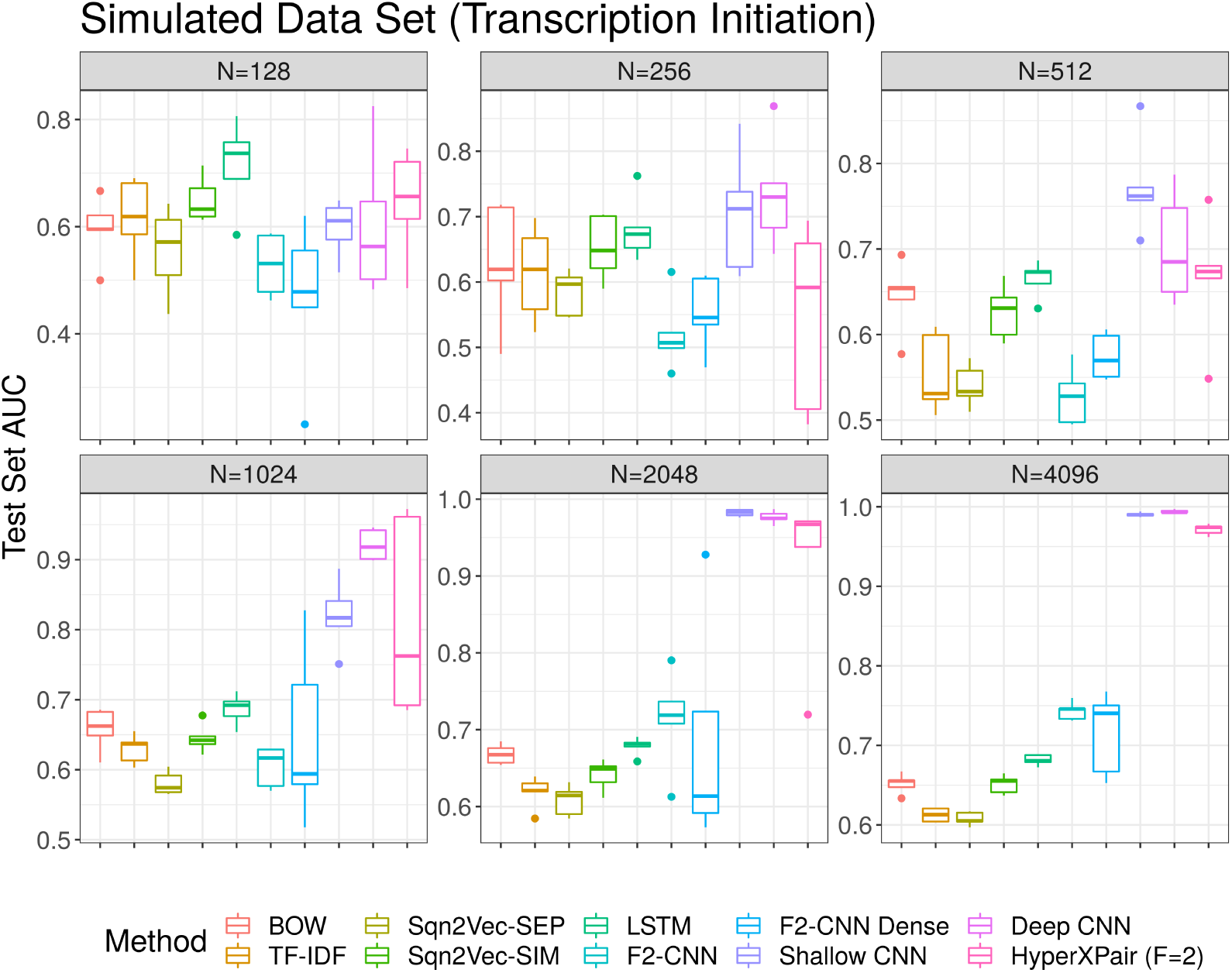
The test set AUC for several methods, as measured on the simulated transcription initiation data (facted by sample size). Our interpretable method, **HyperXPair**, is shown on the far right. Its performance is comparable to LSTM and CNN.

**Figure 5:**
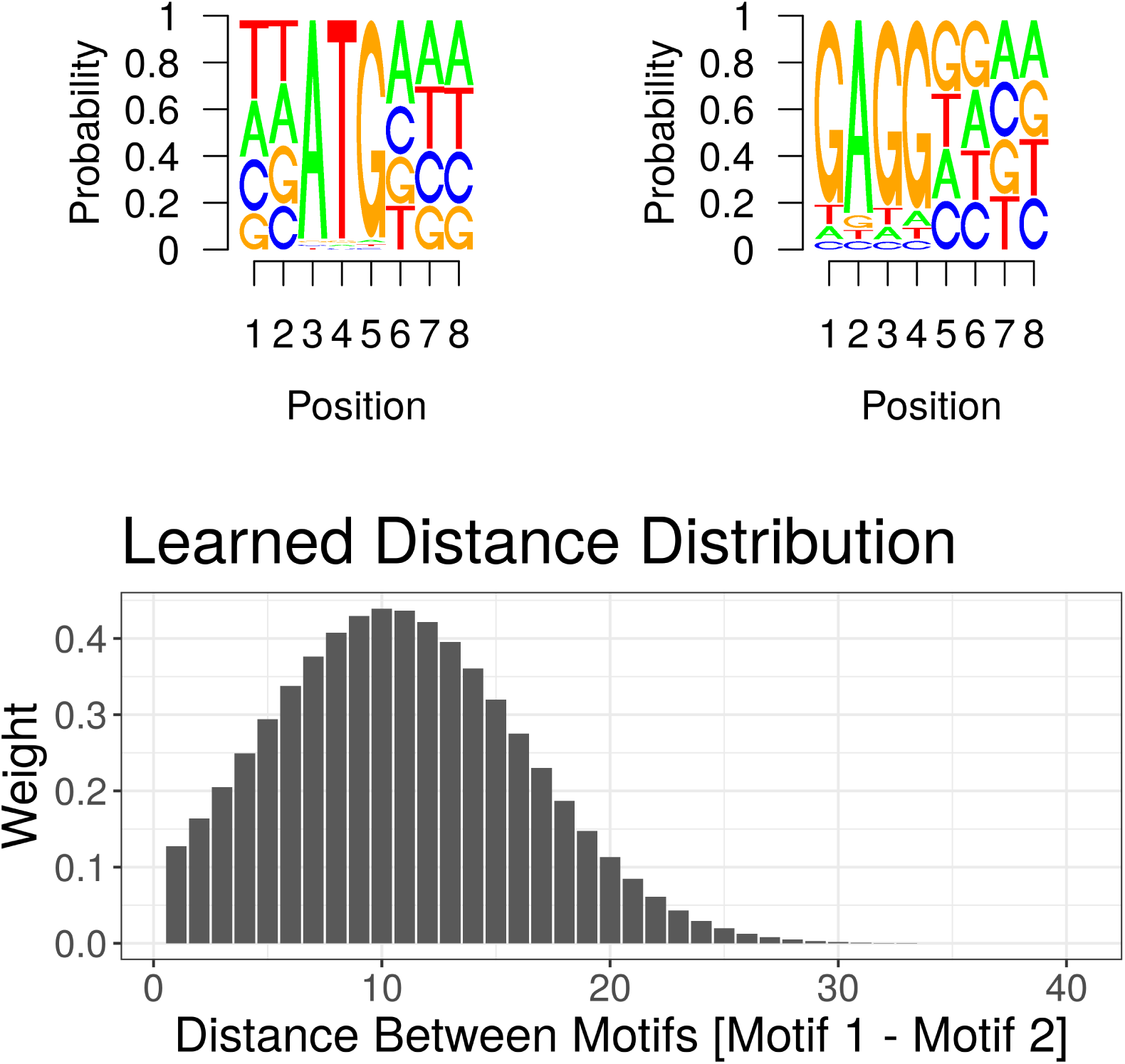
The parameters of the model tell us how it predicts transcription initiation. The top figure shows the softmax-transformed motif weights, which one could convert into regular expressions. For example, the top-left motif would be **ATG***, while the top-right motif would be GAGG****. The bottom panel displays the learned inter-motif distance distribution.

### 6.3 Study 2: Real-world RNA splicing data

We also apply **HyperXPair** to two real data sets, for which we hypothesize motif pair interactions could play a role. In both cases, we want to predict the presence of a splice event in a segment of DNA. The first data set contains labelled donor splice sites (occurring upstream), while the other contains acceptor splice sites (occurring downstream). *Although we hypothesize that motif interactions contribute to splice prediction, we do not think motif interactions define splice prediction*. Nevertheless, these data let us evaluate the real-world utility of **HyperXPair** along two lines of evidence: (1) model performance, and (2) plausibility of learned motifs and distance distributions.

Figure 6 shows the performance of our model and baselines on the donor and acceptor data.

**Figure 6:**
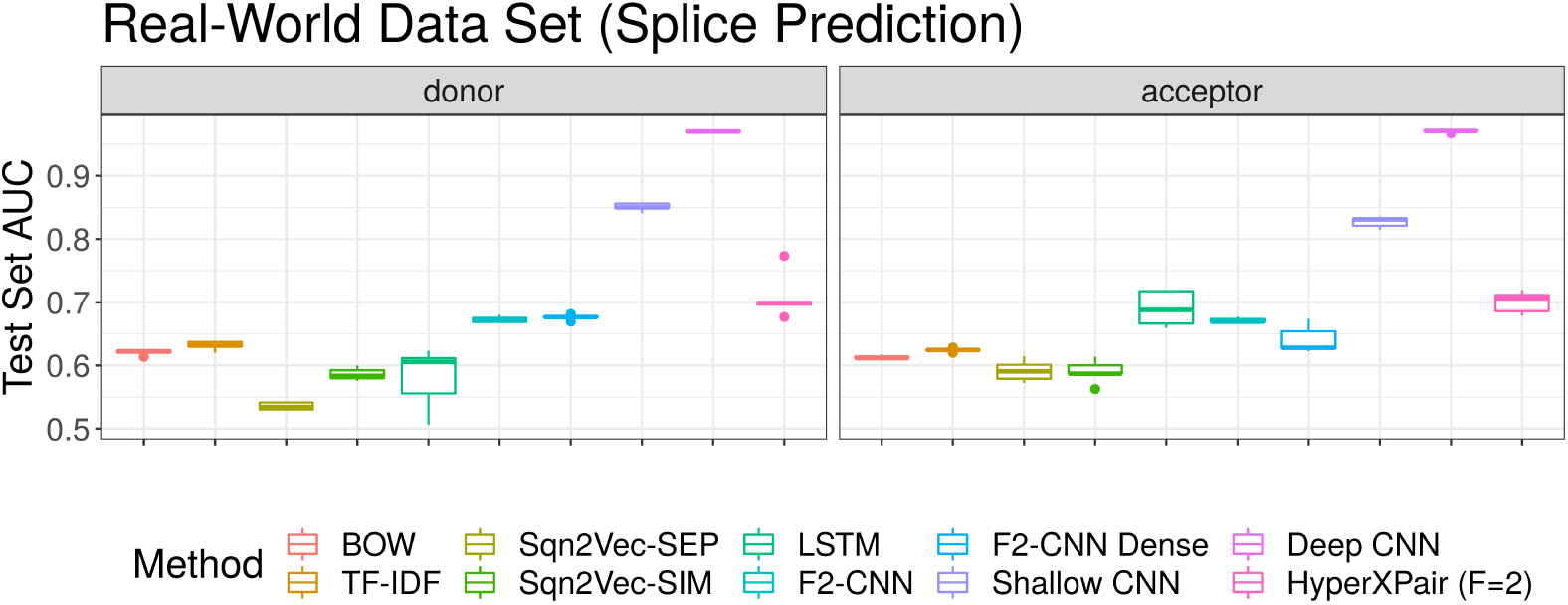
The test set AUC for several methods, as measured on the real-world RNA splicing data (faceted by splice site type). Our interpretable method, **HyperXPair**, is shown on the far right. Its performance is comparable to LSTM, but underperforms the wider and deeper CNN varieties for these data.

Here, we see that running **HyperXPair** with 1 filter pair can achieve an impressive ∼70% AUC, matching the NLP, LSTM, and 2-filter CNN baselines^2^. We also observe, unsurprisingly, that deeper and wider CNNs can outperform **HyperXPair**. This is because our model implies a hypothesis that a single motif pair determines RNA splicing, which is not a complete description of RNA splicing. *However, our primary motivation is not accuracy: rather, it is to obtain an interpretable model of the genomic biology*.

Figure 7A shows the motifs and distances learned for one of the donor training sets. Here, the model learned to associate two motifs, located ∼4 bases apart, with the splice event. In fact, the incredibly low variance of the distance distribution suggests that the model may have learned to stitch together two adjacent motifs into one larger motif. Indeed, we see how (C/A)AGG*T(G/A)AG* from Motif 2 overlaps with *T(G/A)AG*TACC on Motif 1. This finding is important for two reasons: (1) the complete motif almost perfectly resembles the known donor consensus motif ‘(C/A)AGGT(G/A)AG’ to which the U1 small nuclear ribonucleoprotein particle (U1 snRNP) protein binds [Mount et al., 1992], and (2) it shows that **HyperXPair** can still produce meaningful results even when the biological hypothesis about distance-dependent interactions does not apply. Again, this figure explains how the model works, and is produced directly from the model’s parameters.

**Figure 7:**
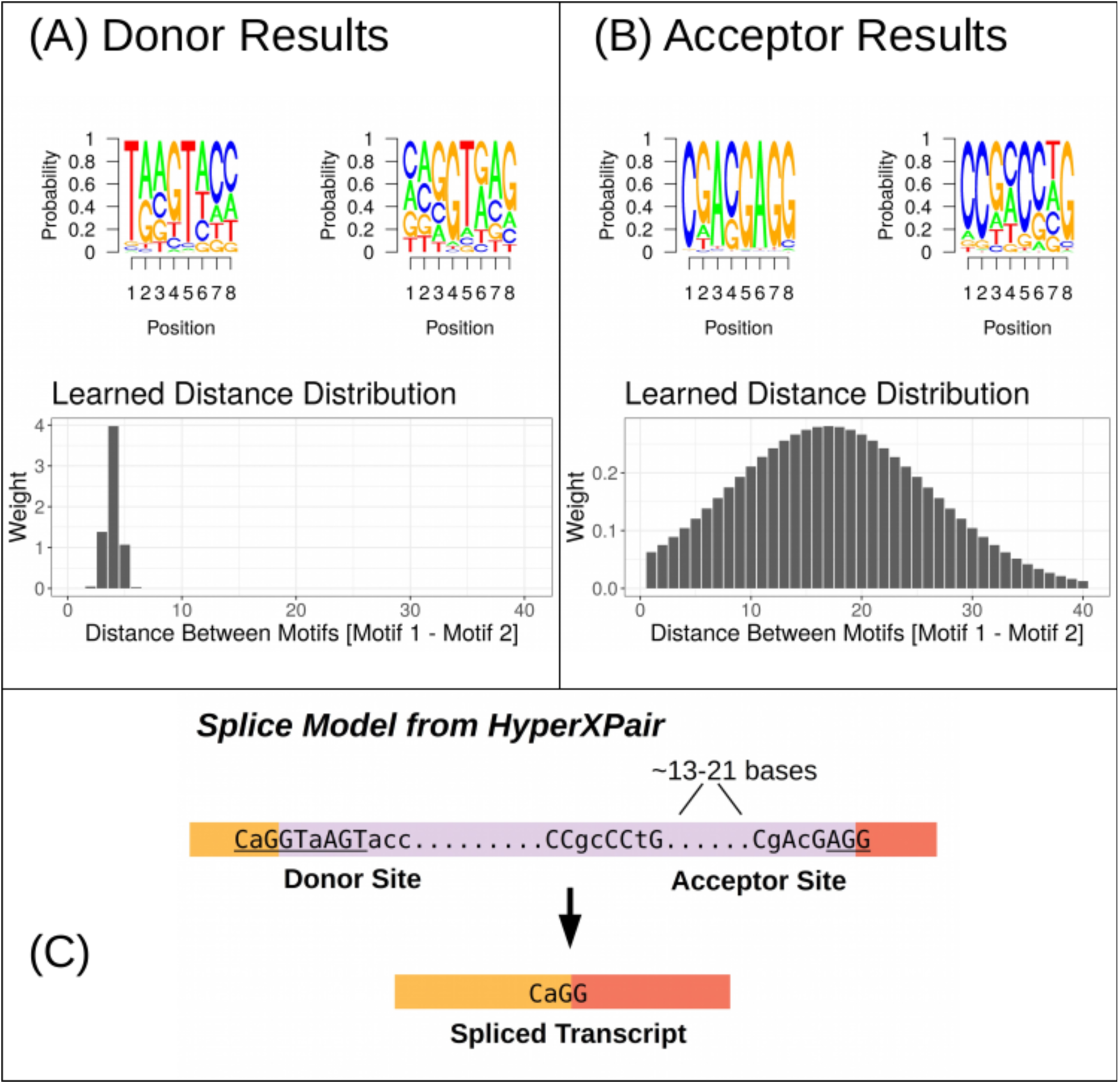
The parameters of the model tell us how it predicts RNA splicing. **Panel A** refers to the **donor** splice site. The model appears to have concatenated two adjacent small motifs – centered around a strong **GT** signal – to form the known donor consensus motif ‘(C/A)AG**GT**(G/A)AG’. **Panel B** refers to the **acceptor** splice site. One motif in the pair appears to contain the canonical **AGG** acceptor splice motif. **Panel C** proposes a schematic of *D. melanogaster* splicing. Lowercase letters are degenerate; underlined letters agree with the literature.

Figure 7B shows the motifs and distances learned for one of the acceptor training sets. Here, the model learned to associate two motifs, located ∼13-21 bases apart, with the splice event. Motif 1 is the downstream motif, which looks like CGACGAGG. This exact 8-base motif shows up in 5/5 training sets, leading us to believe it represents the canonical AGG acceptor splice motif described in the literature [Mount et al., 1992]. Motif 2 is the upstream motif, which looks like CCGCCCTG. This motif shows up in 2/5 training sets, along with the same consensus distance. A look through the literature did not reveal an obvious binding protein for this motif (c.f., Ray et al. [2013]). However, upstream the acceptor site is known to be pyrimidine-rich [Mount et al., 1992], meaning that we should expect Ts or Cs to the *left* of the first motif. The discovered C-rich motif could align with this prior knowledge.

Figure 7C proposes a schematic of *D. melanogaster* splicing based on a synthesis of the **HyperXPair** results with literature (c.f., Mount et al. [1992], Rosenberg et al. [2015]). **Supplemental Figures 10-19** show all filters for the 5 donor and acceptor training sets. **Supplemental Figures 20-29** show all filters for the prior *S* = 160 instead of *S* = 40.

### 6.4 Study 3: Hyper-parameter over-specification

In the real-world setting, we may not know the number of motifs involved. However, **HyperXPair** can still perform well even if we over-specify the number of motifs needed. Using the first *N* = 4096 replicate data set, we repeated the Bayesian optimization scheme for *F* = [3, 4, 5] filters. In all cases, the model successfully learned the GGAGG consensus motif, and achieved an AUC greater than 90%. The model also successfully learned ATG, and the correct GGAGG-ATG distance, when *F* = 4. Interestingly, when *F* = 3 and *F* = 5, the model instead learned the importance of placing GGAGG *after a stop codon* (instead of *before a start codon*). Although this is not the exact rule we expected **HyperXPair** to learn, it is a valid rule given the simulated data. (It also highlights how filters can learn degenerate motifs: TAG, TAA, and TGA are all stop codons, 2 of which get captured by the “T(A/G)A” filter.)

Note that the appropriate motifs emerge without any regularization on the CNN filters themselves. The only regularization in our model occurs at the final layer, which is a simple regression. We expect that aggressively regularizing the CNN filters, for example as proposed by [Koo et al., 2019] and [Ploenzke and Irizarry, 2018], could further improve the motif identity module when needed. **Supplemental Figures 7-9** show all filters for the *F* = [3, 4, 5] runs.

## 7 Conclusion

Although a deep CNN can implicitly learn motif interactions, the identity of the motifs, and the relevant distances between them, are not readily interpretable from the model. In contrast, our proposed neural network architecture can answer two questions: (1) What are the consensus motifs? and (2) What are the inter-motif distance distributions?

By learning the motifs and distance distributions explicitly, **HyperXPair** can offer insights into the mechaninistic basis of the genomic event under study. In the case of transcription initiation, we recovered knowledge that the Shine-Dalgarno sequence should occur ∼5-13 bases upstream from the Kozak consensus sequence. In the case of *D. melanogaster* splicing, we verified established knowledge about the donor and acceptor splice sites, and discovered a novel motif pair: a C-rich motif occurring ∼13-21 bases upstream from the canonical acceptor splice site. These findings demonstrate how neural networks can be adapted to advance knowledge in genomic biology.

## 8 Data Availability

Code and simulated data will be publicly available after peer review. Splice data are available from https://github.com/SomayahAlbaradei/Splice_Deep.

## Footnotes

1 https://github.com/SomayahAlbaradei/Splice_Deep

2 A comment on the AUC reported by Albaradei et al. [2020], and how it relates to both global pooling and the AUC reported here, is provided in the **Supplement**.

## References

Sophie C. Bonnal, Irene López-Oreja, and Juan Valcárcel. Roles and mechanisms of alternative splicing in cancer - implications for care. Nature Reviews. Clinical Oncology, 17(8):457–474, August 2020. ISSN 1759-4782. doi: 10.1038/s41571-020-0350-x.

Travers Ching, Daniel S. Himmelstein, Brett K. Beaulieu-Jones, Alexandr A. Kalinin, Brian T. Do, Gregory P. Way, Enrico Ferrero, Paul-Michael Agapow, Michael Zietz, Michael M. Hoffman, Wei Xie, Gail L. Rosen, Benjamin J. Lengerich, Johnny Israeli, Jack Lanchantin, Stephen Woloszynek, Anne E. Carpenter, Avanti Shrikumar, Jinbo Xu, Evan M. Cofer, Christopher A. Lavender, Srinivas C. Turaga, Amr M. Alexandari, Zhiyong Lu, David J. Harris, Dave DeCaprio, Yanjun Qi, Anshul Kundaje, Yifan Peng, Laura K. Wiley, Marwin H. S. Segler, Simina M. Boca, S. Joshua Swamidass, Austin Huang, Anthony Gitter, and Casey S. Greene. Opportunities and obstacles for deep learning in biology and medicine. Journal of the Royal Society, Interface, 15 (141), 2018. ISSN 1742-5662. doi: 10.1098/rsif.2017.0387.

Kishore Jaganathan, Sofia Kyriazopoulou Panagiotopoulou, Jeremy F. McRae, Siavash Fazel Darbandi, David Knowles, Yang I. Li, Jack A. Kosmicki, Juan Arbelaez, Wenwu Cui, Grace B. Schwartz, Eric D. Chow, Efstathios Kanterakis, Hong Gao, Amirali Kia, Serafim Batzoglou, Stephan J. Sanders, and Kyle Kai-How Farh. Predicting Splicing from Primary Sequence with Deep Learning. Cell, 176(3):535–548.e24, January 2019. ISSN 0092-8674. doi: 10.1016/j.cell.2018.12.015. URL http://www.sciencedirect.com/science/article/pii/S0092867418316295.

Ruohan Wang, Zishuai Wang, Jianping Wang, and Shuaicheng Li. SpliceFinder: ab initio prediction of splice sites using convolutional neural network. BMC Bioinformatics, 20(23):652, December 2019. ISSN 1471-2105. doi: 10.1186/s12859-019-3306-3. URL https://doi.org/10.1186/s12859-019-3306-3.

Somayah Albaradei, Arturo Magana-Mora, Maha Thafar, Mahmut Uludag, Vladimir B. Bajic, Takashi Gojobori, Magbubah Essack, and Boris R. Jankovic. Splice2Deep: An ensemble of deep convolutional neural networks for improved splice site prediction in genomic DNA. Gene: X, 5:100035, December 2020. ISSN 2590-1583. doi: 10.1016/j.gene.2020.100035. URL http://www.sciencedirect.com/science/article/pii/S2590158320300097.

Jiong Ma, Allan Campbell, and Samuel Karlin. Correlations between Shine-Dalgarno Sequences and Gene Features Such as Predicted Expression Levels and Operon Structures. Journal of Bacteriology, 184(20):5733–5745, October 2002. ISSN 0021-9193. doi: 10.1128/JB.184.20.5733-5745.2002. URL https://www.ncbi.nlm.nih.gov/pmc/articles/PMC139613/.

Shengdong Ke and Lawrence Allen Chasin. Intronic motif pairs cooperate across exons to promote pre-mRNA splicing. 11(R84), 2010. doi: 10.7916/D84F1P46. URL https://doi.org/10.7916/D84F1P46.

Bobak Shahriari, Kevin Swersky, Ziyu Wang, Ryan P. Adams, and Nando de Freitas. Taking the Human Out of the Loop: A Review of Bayesian Optimization. Proceedings of the IEEE, 104(1): 148–175, January 2016. ISSN 1558-2256. doi: 10.1109/JPROC.2015.2494218.

Jernej Ule and Benjamin J. Blencowe. Alternative Splicing Regulatory Networks: Functions, Mechanisms, and Evolution. Molecular Cell, 76(2):329–345, October 2019. ISSN 1097-2765. doi: 10.1016/j.molcel.2019.09.017. URL http://www.sciencedirect.com/science/article/pii/S1097276519307026.

Ka-Chun Wong, Yue Li, and Chengbin Peng. Identification of coupling DNA motif pairs on long-range chromatin interactions in human K562 cells. Bioinformatics, 32(3):321–324, February 2016. ISSN 1367-4803. doi: 10.1093/bioinformatics/btv555. URL https://academic.oup.com/bioinformatics/article/32/3/321/1743368.

Jian Zhou and Olga G. Troyanskaya. Predicting effects of noncoding variants with deep learning–based sequence model. Nature Methods, 12(10):931–934, October 2015. ISSN 1548-7105. doi: 10.1038/nmeth.3547. URL https://www.nature.com/articles/nmeth.3547.

Peter K. Koo, Praveen Anand, Steffan B. Paul, and Sean R. Eddy. Inferring Sequence-Structure Preferences of RNA-Binding Proteins with Convolutional Residual Networks. bioRxiv, page 418459, September 2018. doi: 10.1101/418459. URL https://www.biorxiv.org/content/10.1101/418459v1.

Avanti Shrikumar, Peyton Greenside, and Anshul Kundaje. Learning Important Features Through Propagating Activation Differences. arXiv:1704.02685 [cs], October 2019. URL http://arxiv.org/abs/1704.02685. arXiv: 1704.02685.

David Alvarez-Melis and Tommi S. Jaakkola. Towards Robust Interpretability with Self-Explaining Neural Networks. June 2018. URL https://arxiv.org/abs/1806.07538v2.

John Hawkins and Mikael Bodén. The Applicability of Recurrent Neural Networks for Biological Sequence Analysis. IEEE/ACM Transactions on Computational Biology and Bioinformatics, 2: 243–253, 2005.

Sepp Hochreiter and Jürgen Schmidhuber. Long Short-Term Memory. Neural Computation, 9(8): 1735–1780, November 1997. ISSN 0899-7667. doi: 10.1162/neco.1997.9.8.1735. URL https://doi.org/10.1162/neco.1997.9.8.1735.

Hamid Reza Hassanzadeh and May D. Wang. DeeperBind: Enhancing Prediction of Sequence Specificities of DNA Binding Proteins. November 2016. URL https://arxiv.org/abs/1611.05777v1.

Xiaoyong Pan, Peter Rijnbeek, Junchi Yan, and Hong-Bin Shen. Prediction of RNA-protein sequence and structure binding preferences using deep convolutional and recurrent neural networks. BMC Genomics, 19(1):511, July 2018. ISSN 1471-2164. doi: 10.1186/s12864-018-4889-1. URLhttps://doi.org/10.1186/s12864-018-4889-1.

Kunihiko Fukushima. Neocognitron: A self-organizing neural network model for a mechanism of pattern recognition unaffected by shift in position. Biological Cybernetics, 36(4):193–202, April 1980. ISSN 1432-0770. doi: 10.1007/BF00344251. URL https://doi.org/10.1007/BF00344251.

A. Waibel, T. Hanazawa, G. Hinton, K. Shikano, and K.J. Lang. Phoneme recognition using time-delay neural networks. IEEE Transactions on Acoustics, Speech, and Signal Processing, 37(3): 328–339, March 1989. ISSN 0096-3518. doi: 10.1109/29.21701.

Thin Nguyen, Hang Le, Thomas P. Quinn, Thuc Le, and Svetha Venkatesh. GraphDTA: Predicting drug–target binding affinity with graph neural networks. bioRxiv, page 684662, April 2020a. doi: 10.1101/684662. URL https://www.biorxiv.org/content/10.1101/684662v7.

Suying Bao, Daniel F. Moakley, and Chaolin Zhang. The Splicing Code Goes Deep. Cell, 176(3): 414–416, January 2019. ISSN 0092-8674. doi: 10.1016/j.cell.2019.01.013. URL http://www.sciencedirect.com/science/article/pii/S0092867419300467.

Peter K. Koo and Sean R. Eddy. Representation learning of genomic sequence motifs with convolutional neural networks. PLOS Computational Biology, 15(12):e1007560, December 2019. ISSN 1553-7358. doi: 10.1371/journal.pcbi.1007560. URL https://journals.plos.org/ploscompbiol/article?id=10.1371/journal.pcbi.1007560.

Peter K. Koo and Matt Ploenzke. Improving Convolutional Network Interpretability with Exponential Activations. bioRxiv, page 650804, May 2019. doi: 10.1101/650804. URL https://www.biorxiv.org/content/10.1101/650804v1.

Peter K. Koo, Sharon Qian, Gal Kaplun, Verena Volf, and Dimitris Kalimeris. Robust Neural Networks are More Interpretable for Genomics. bioRxiv, page 657437, June 2019. doi: 10.1101/657437. URL https://www.biorxiv.org/content/10.1101/657437v1.

M. S. Ploenzke and R. A. Irizarry. Interpretable Convolution Methods for Learning Genomic Sequence Motifs. bioRxiv, page 411934, September 2018. doi: 10.1101/411934. URL https://www.biorxiv.org/content/10.1101/411934v1.

T D Schneider and R M Stephens. Sequence logos: a new way to display consensus sequences. Nucleic Acids Research, 18(20):6097–6100, October 1990. ISSN 0305-1048. URL https://www.ncbi.nlm.nih.gov/pmc/articles/PMC332411/.

Jasper Snoek, Hugo Larochelle, and Ryan Adams. Practical bayesian optimization of machine learning algorithms. In NIPS, pages 2951–2959, 2012.

Dang Nguyen, Sunil Gupta, Santu Rana, Alistair Shilton, and Svetha Venkatesh. Bayesian optimization for categorical and category-specific continuous inputs. In AAAI, 2020b.

Carl Rasmussen. Gaussian processes in machine learning. In Summer School on Machine Learning, pages 63–71. Springer, 2003.

Niranjan Srinivas, Andreas Krause, Sham Kakade, and Matthias Seeger. Information-theoretic regret bounds for gaussian process optimization in the bandit setting. IEEE Transactions on Information Theory, 58(5):3250–3265, 2012.

G D Stormo, T D Schneider, L Gold, and A Ehrenfeucht. Use of the ‘Perceptron’ algorithm to distinguish translational initiation sites in E. coli. Nucleic Acids Research, 10(9):2997–3011, May 1982. ISSN 0305-1048. URL https://www.ncbi.nlm.nih.gov/pmc/articles/PMC320670/.

S M Mount, C Burks, G Hertz, G D Stormo, O White, and C Fields. Splicing signals in Drosophila: intron size, information content, and consensus sequences. Nucleic Acids Research, 20(16):4255– 4262, August 1992. ISSN 0305-1048. URL https://www.ncbi.nlm.nih.gov/pmc/articles/PMC334133/.

Sanjeev Arora, Mikhail Khodak, Nikunj Saunshi, and Kiran Vodrahalli. A Compressed Sensing View of Unsupervised Text Embeddings, Bag-of-n-Grams, and LSTMs. February 2018. URL https://openreview.net/forum?id=B1e5ef-C-.

Dang Nguyen, Wei Luo, Tu Dinh Nguyen, Svetha Venkatesh, and Dinh Phung. Sqn2vec: Learning sequence representation via sequential patterns with a gap constraint. In ECML-PKDD, pages 569–584. Springer, 2018.

Debashish Ray, Hilal Kazan, Kate B. Cook, Matthew T. Weirauch, Hamed S. Najafabadi, Xiao Li, Serge Gueroussov, Mihai Albu, Hong Zheng, Ally Yang, Hong Na, Manuel Irimia, Leah H. Matzat, Ryan K. Dale, Sarah A. Smith, Christopher A. Yarosh, Seth M. Kelly, Behnam Nabet, Desirea Mecenas, Weimin Li, Rakesh S. Laishram, Mei Qiao, Howard D. Lipshitz, Fabio Piano, Anita H. Corbett, Russ P. Carstens, Brendan J. Frey, Richard A. Anderson, Kristen W. Lynch, Luiz O. F. Penalva, Elissa P. Lei, Andrew G. Fraser, Benjamin J. Blencowe, Quaid D. Morris, and Timothy R. Hughes. A compendium of RNA-binding motifs for decoding gene regulation. Nature, 499(7457):172–177, July 2013. ISSN 0028-0836. doi: 10.1038/nature12311. URL https://www.ncbi.nlm.nih.gov/pmc/articles/PMC3929597/.

Alexander B. Rosenberg, Rupali P. Patwardhan, Jay Shendure, and Georg Seelig. Learning the sequence determinants of alternative splicing from millions of random sequences. Cell, 163(3): 698–711, October 2015. ISSN 1097-4172. doi: 10.1016/j.cell.2015.09.054.

