## Supplement for "Learning Distance-Dependent Motif Interactions: An Explicitly Interpretable Neural Model of Genomic Events"

#### 1 A comment on global pooling

We should comment on the apparent discrepancy between the accuracies from our model and those from Albaradei et al. [2020], who reported >98.8% AUC for simple CNNs. We hypothesize that this follows from an artifact in the training and test data that allows positional information to leak through. In both the training and test data, the splice event always occurs at position 301. Without global pooling, a CNN can keep positional information intact, allowing the model to learn that only bases surrounding position 301 matter. During testing, the model could guess most samples correctly just by checking, for example, if an acceptor motif like AGG occurs at position 301. This kind of model will perform well during testing, but would fail in a real-world scenario, even where the acceptor site could occur anywhere. Indeed, a simple motif like AGG would occur throughout the genome where splicing does not occur. Without global pooling, we can achieve 97% AUC with a single CNN filter, but this is not a generally useful model. In contrast, **HyperXPair** uses global pooling to erase positional information; although this appears to reduce accuracy, it actually yields a more robust model of RNA splicing.

#### References


Somayah Albaradei, Arturo Magana-Mora, Maha Thafar, Mahmut Uludag, Vladimir B. Bajic, Takashi Gojobori, Magbubah Essack, and Boris R. Jankovic. Splice2Deep: An ensemble of deep convolutional neural networks for improved splice site prediction in genomic DNA. *Gene: X*, 5:100035, December 2020. ISSN 2590-1583. doi: 10.1016/j.gene.2020.100035. URL http://www.sciencedirect.com/science/article/pii/S2590158320300097.

#### 2 Overview

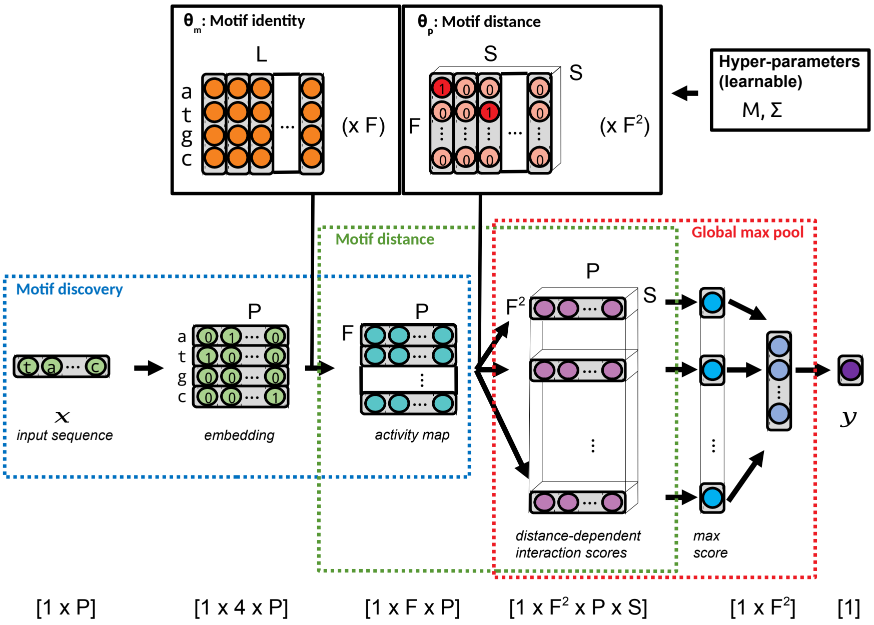


Figure 1: This figure shows an overview of the **HyperXPair** architecture. First, a CNN learns motif identities from a 1-hot embedding of DNA, producing an **activity map** that describes the presence of each motif along the sequence. Second, another CNN uses a motif co-occurrence kernel to compute co-occurrence scores for each motif pair along the sequence. Third, a global max pool layer selects the largest co-occurrence score for each pair, which are then used to predict the outcome via a simple regression. Note that the tensor sizes shown in the schematic are simplified for visualization, and do not include padding.

### 3 Simulated Data Results (F=2)

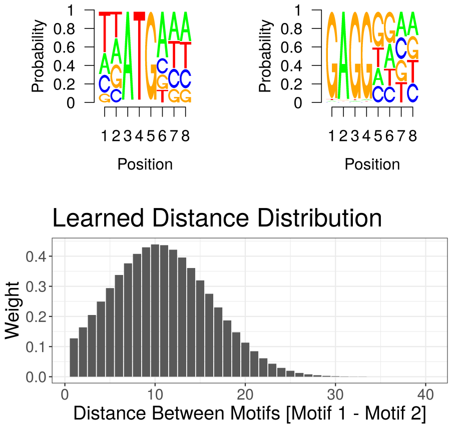


Figure 2: This figure shows the raw filter weights, corresponding seqLogo plots, and inter-motif distances for a single filter pair, used to predict **transcription initiation**. All relevant parameters and hyper-parameters were learned from the first $N = 4096$ replicate training set.

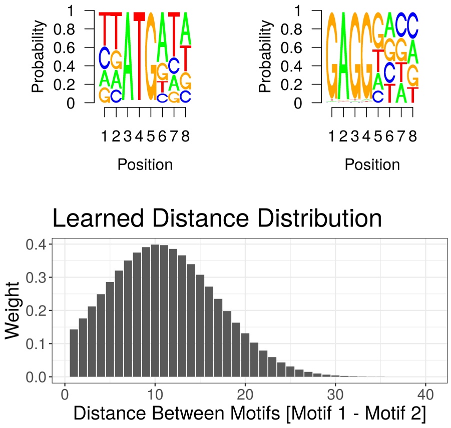


Figure 3: This figure shows the raw filter weights, corresponding seqLogo plots, and inter-motif distances for a single filter pair, used to predict **transcription initiation**. All relevant parameters and hyper-parameters were learned from the second $N = 4096$ replicate training set.

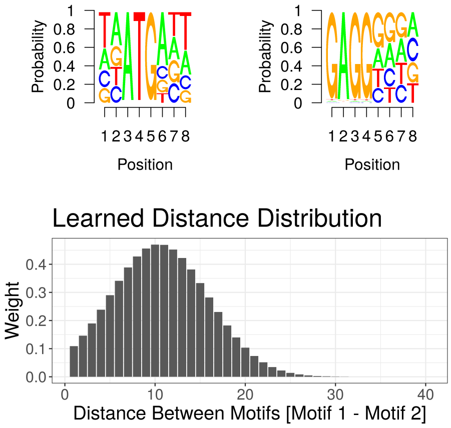


Figure 4: This figure shows the raw filter weights, corresponding seqLogo plots, and inter-motif distances for a single filter pair, used to predict **transcription initiation**. All relevant parameters and hyper-parameters were learned from the third $N = 4096$ replicate training set.

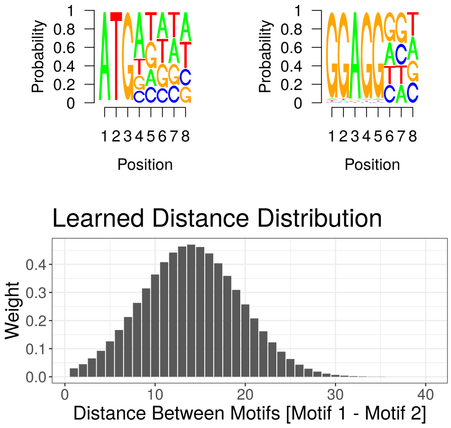


Figure 5: This figure shows the raw filter weights, corresponding seqLogo plots, and inter-motif distances for a single filter pair, used to predict **transcription initiation**. All relevant parameters and hyper-parameters were learned from the fourth $N = 4096$ replicate training set.

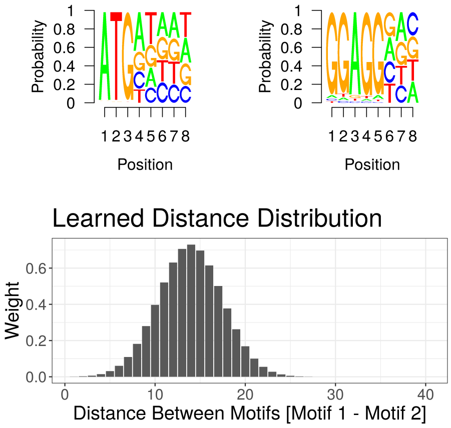


Figure 6: This figure shows the raw filter weights, corresponding seqLogo plots, and inter-motif distances for a single filter pair, used to predict **transcription initiation**. All relevant parameters and hyper-parameters were learned from the fifth $N = 4096$ replicate training set.

### 4 Simulated Data Results (F>2)

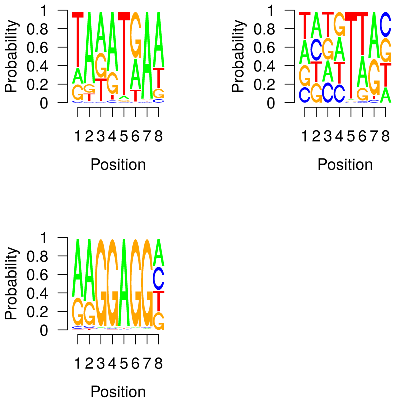


Figure 7: This figure shows the raw filter weights and corresponding seqLogo plots for all filter pairs, used to predict **transcription initiation**. All relevant parameters and hyper-parameters were learned from the first $N = 4096$ replicate training set using **3 total filters**.

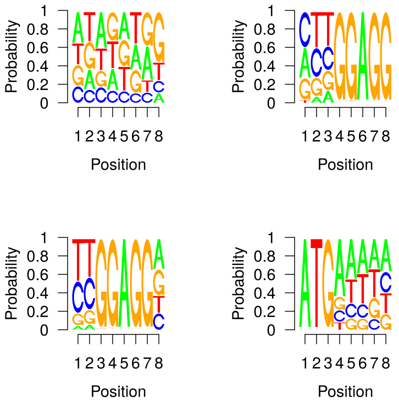


Figure 8: This figure shows the raw filter weights and corresponding seqLogo plots for all filter pairs, used to predict **transcription initiation**. All relevant parameters and hyper-parameters were learned from the first $N = 4096$ replicate training set using **4 total filters**.

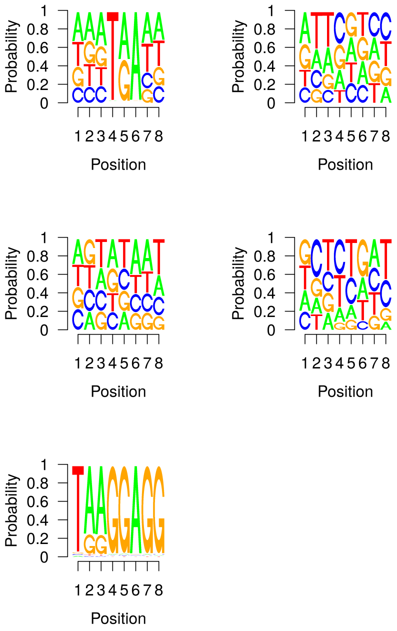


Figure 9: This figure shows the raw filter weights and corresponding seqLogo plots for all filter pairs, used to predict **transcription initiation**. All relevant parameters and hyper-parameters were learned from the first $N = 4096$ replicate training set using **5 total filters**.

### 5 Fruit Fly Donor Results (F=2; S=40)

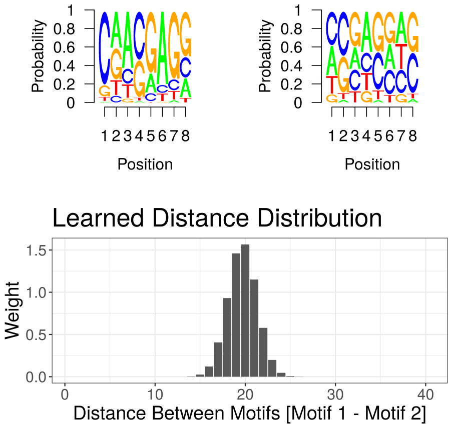


Figure 10: This figure shows the raw filter weights, corresponding seqLogo plots, and inter-motif distances for a single filter pair, used to predict a **donor splice event**. All relevant parameters and hyper-parameters were learned from the first donor training set.

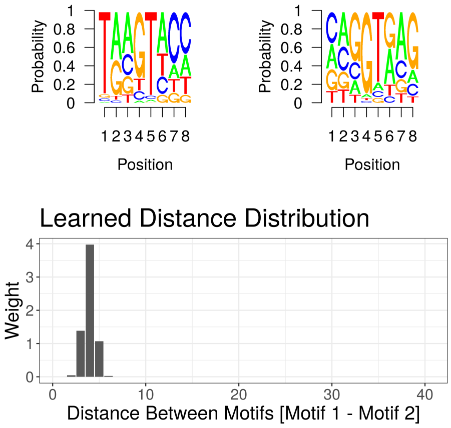


Figure 11: This figure shows the raw filter weights, corresponding seqLogo plots, and inter-motif distances for a single filter pair, used to predict a **donor splice event**. All relevant parameters and hyper-parameters were learned from the second donor training set.

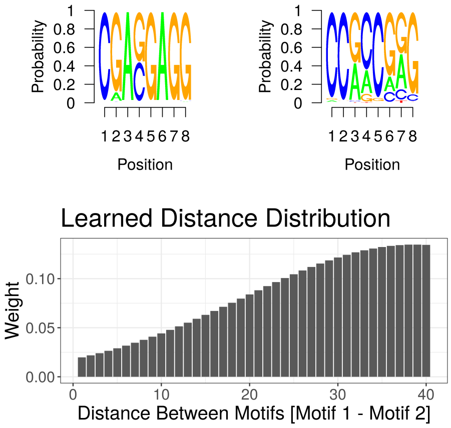


Figure 12: This figure shows the raw filter weights, corresponding seqLogo plots, and inter-motif distances for a single filter pair, used to predict a **donor splice event**. All relevant parameters and hyper-parameters were learned from the third donor training set.

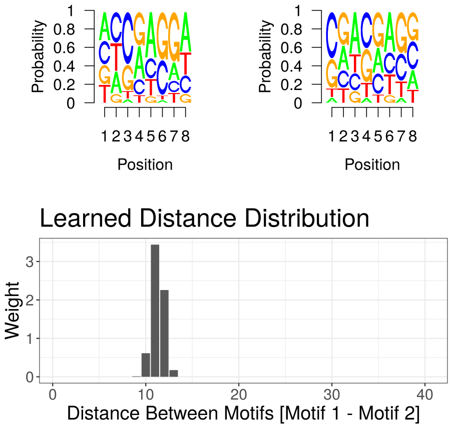


Figure 13: This figure shows the raw filter weights, corresponding seqLogo plots, and inter-motif distances for a single filter pair, used to predict a **donor splice event**. All relevant parameters and hyper-parameters were learned from the fourth donor training set.

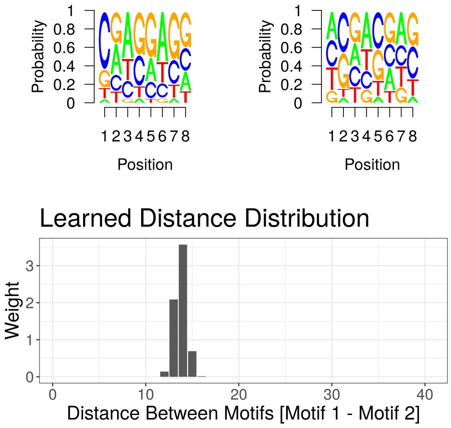


Figure 14: This figure shows the raw filter weights, corresponding seqLogo plots, and inter-motif distances for a single filter pair, used to predict a **donor splice event**. All relevant parameters and hyper-parameters were learned from the fifth donor training set.

### 6 Fruit Fly Acceptor Results (F=2; S=40)

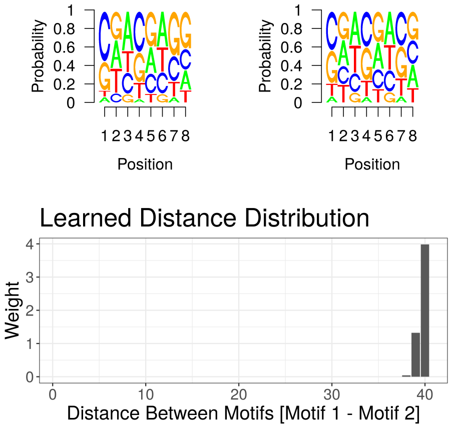


Figure 15: This figure shows the raw filter weights, corresponding seqLogo plots, and inter-motif distances for a single filter pair, used to predict a **acceptor splice event**. All relevant parameters and hyper-parameters were learned from the first acceptor training set.

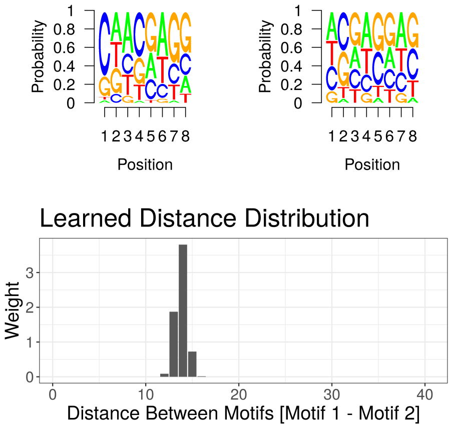


Figure 16: This figure shows the raw filter weights, corresponding seqLogo plots, and inter-motif distances for a single filter pair, used to predict a **acceptor splice event**. All relevant parameters and hyper-parameters were learned from the second acceptor training set.

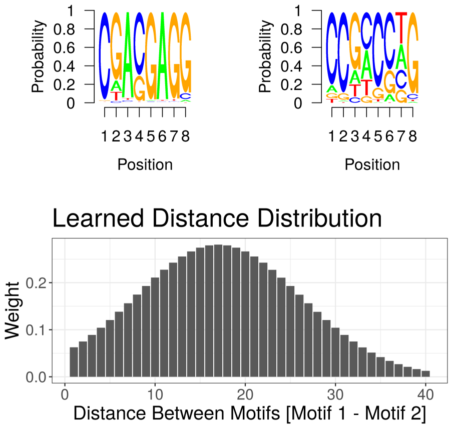


Figure 17: This figure shows the raw filter weights, corresponding seqLogo plots, and inter-motif distances for a single filter pair, used to predict a **acceptor splice event**. All relevant parameters and hyper-parameters were learned from the third acceptor training set.

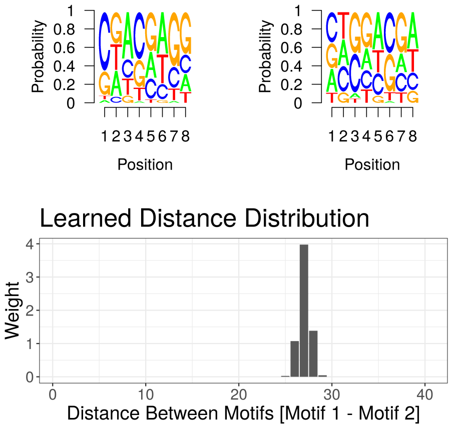


Figure 18: This figure shows the raw filter weights, corresponding seqLogo plots, and inter-motif distances for a single filter pair, used to predict a **acceptor splice event**. All relevant parameters and hyper-parameters were learned from the fourth acceptor training set.

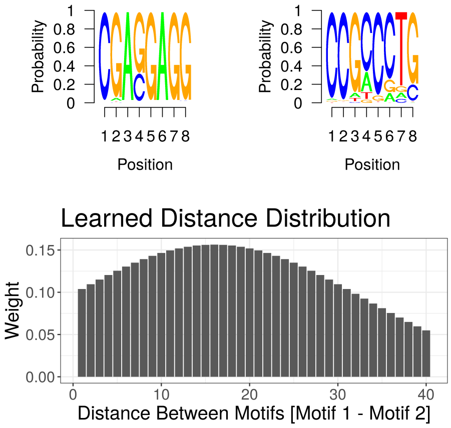


Figure 19: This figure shows the raw filter weights, corresponding seqLogo plots, and inter-motif distances for a single filter pair, used to predict a **acceptor splice event**. All relevant parameters and hyper-parameters were learned from the fifth acceptor training set.

### 7 Fruit Fly Donor Results (F=2; S=160)

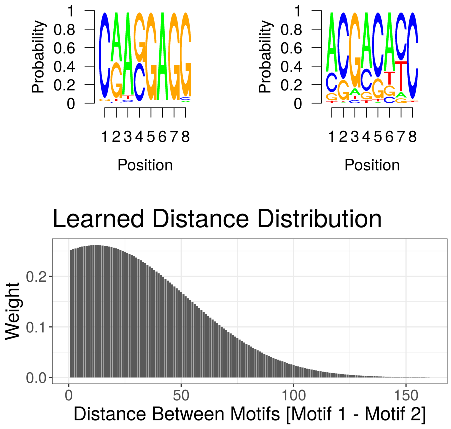


Figure 20: This figure shows the raw filter weights, corresponding seqLogo plots, and inter-motif distances for a single filter pair, used to predict a **donor splice event**. All relevant parameters and hyper-parameters were learned from the first donor training set.

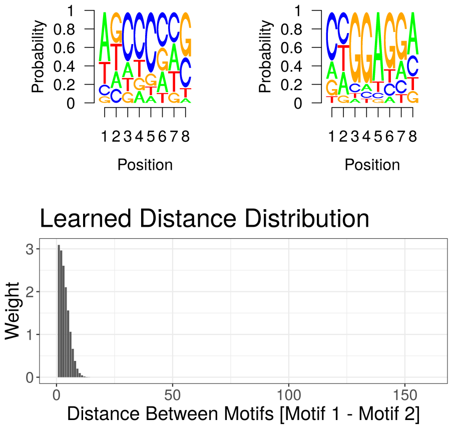


Figure 21: This figure shows the raw filter weights, corresponding seqLogo plots, and inter-motif distances for a single filter pair, used to predict a **donor splice event**. All relevant parameters and hyper-parameters were learned from the second donor training set.

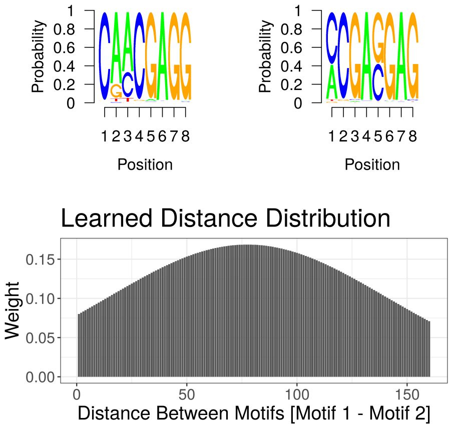


Figure 22: This figure shows the raw filter weights, corresponding seqLogo plots, and inter-motif distances for a single filter pair, used to predict a **donor splice event**. All relevant parameters and hyper-parameters were learned from the third donor training set.

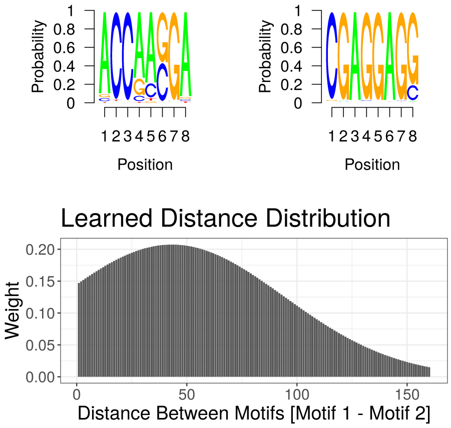


Figure 23: This figure shows the raw filter weights, corresponding seqLogo plots, and inter-motif distances for a single filter pair, used to predict a **donor splice event**. All relevant parameters and hyper-parameters were learned from the fourth donor training set.

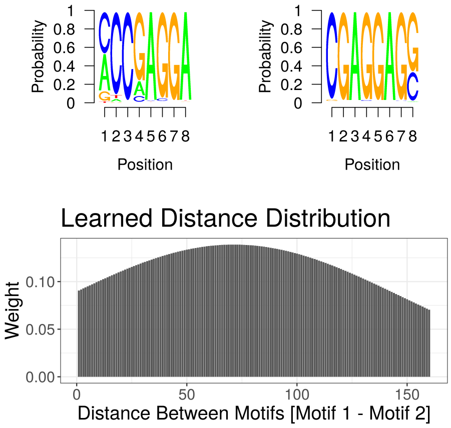


Figure 24: This figure shows the raw filter weights, corresponding seqLogo plots, and inter-motif distances for a single filter pair, used to predict a **donor splice event**. All relevant parameters and hyper-parameters were learned from the fifth donor training set.

### 8 Fruit Fly Acceptor Results (F=2; S=160)

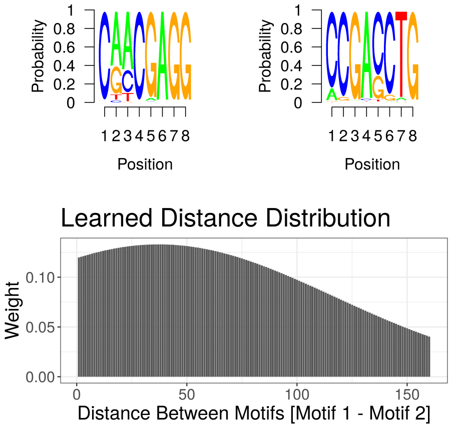


Figure 25: This figure shows the raw filter weights, corresponding seqLogo plots, and inter-motif distances for a single filter pair, used to predict a **acceptor splice event**. All relevant parameters and hyper-parameters were learned from the first acceptor training set.

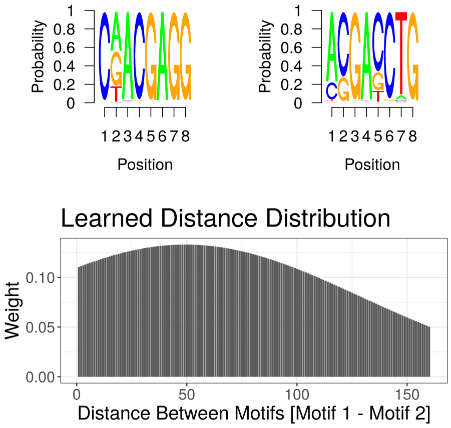


Figure 26: This figure shows the raw filter weights, corresponding seqLogo plots, and inter-motif distances for a single filter pair, used to predict a **acceptor splice event**. All relevant parameters and hyper-parameters were learned from the second acceptor training set.

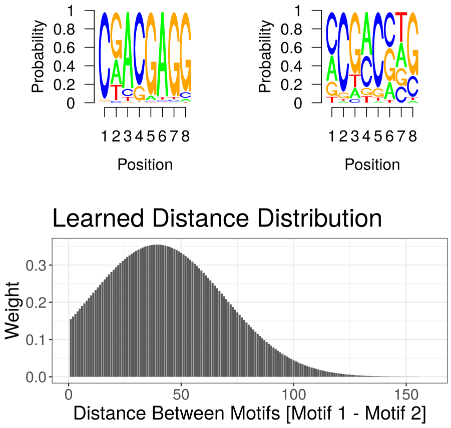


Figure 27: This figure shows the raw filter weights, corresponding seqLogo plots, and inter-motif distances for a single filter pair, used to predict a **acceptor splice event**. All relevant parameters and hyper-parameters were learned from the third acceptor training set.

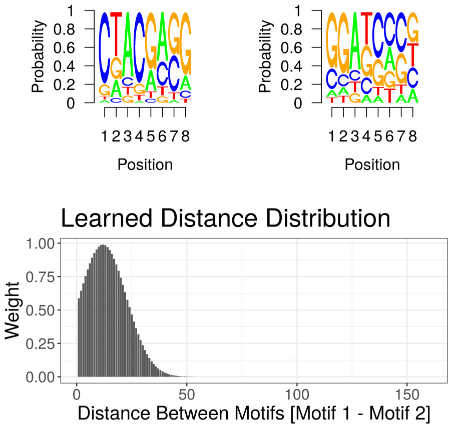


Figure 28: This figure shows the raw filter weights, corresponding seqLogo plots, and inter-motif distances for a single filter pair, used to predict a **acceptor splice event**. All relevant parameters and hyper-parameters were learned from the fourth acceptor training set.

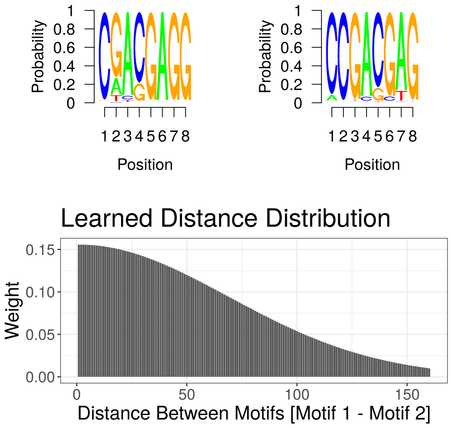


Figure 29: This figure shows the raw filter weights, corresponding seqLogo plots, and inter-motif distances for a single filter pair, used to predict a **acceptor splice event**. All relevant parameters and hyper-parameters were learned from the fifth acceptor training set.
